# Biochemistry and nanomechanical properties of human colon cells upon simvastatin, lovastatin and mevastatin supplementations – Raman imaging and AFM studies

**DOI:** 10.1101/2022.01.19.476934

**Authors:** K. Beton, B. Brożek-Płuska

**Affiliations:** Lodz University of Technology, Institute of Applied Radiation Chemistry, Laboratory of Laser Molecular Spectroscopy, Wroblewskiego 15, 93-590 Lodz and Poland

## Abstract

One of the most important areas of medical science is oncology, which is responsible for both the diagnostics and treatment of cancer diseases. Over the years, there has been an intensive development of cancer diagnostics and treatment. This paper shows the comparison of normal (CCD-18Co) and cancerous (CaCo-2) cell lines of the human gastrointestinal tract on the basis of nanomechanical and biochemical properties in order to obtain information on the cancer biomarkers useful in oncological diagnostics. The research techniques used were: Atomic Force Microscopy and Raman spectroscopy and imaging. In addition, the studies included also the effect of the statins compounds: mevastatin, lovastatin and simvastatin and their influence on nanomechanical and biochemical changes of properties of cells tracking using AFM and Raman imaging techniques. The cytotoxicity of mevastatin and lovastatin was determined by using XTT tests.

## Introduction

Cancer development is a complex multi-stage process related to the transformations of normal cells to pathological. Colorectal cancer (CRC) is the second most common cancer both in men and woman worldwide, and is the leading cause of death. The mortality rate related to this type of cancer is high and approximately equal to 60% in Europe and USA [1]. Moreover, CRC is characterized with high metastasis [2–4]. The risk factors for CRC can be divided into three main groups: 1 environmental (e.g. high-fat diet, high-calorie diet, diet low in silage, vegetables and fruit), 2 internal (e.g. adenomas, ulcers, Crohn’s syndrome), and 3 genetic (e.g. familial adenomatous polyposis) [5]. 75–95% of CRC cases occur in people without any genetic load, which makes lifestyle and eating habits particularly important in this type of cancer development [6,7].

Generally, in the first stage of CRC development healthy cells in the lining of the colon change, grow, and divide uncontrollably to form a mass of tumor. Both genetic and environmental factors can change the dynamics of this process. CRC most often begins with a polyp, a non-cancerous growth that can develop on the inner wall of the colon and then can transform into cancer or metastatic cancer. Scheme 1 presents the cross section through the layers of the human colon.

**Scheme 1.**
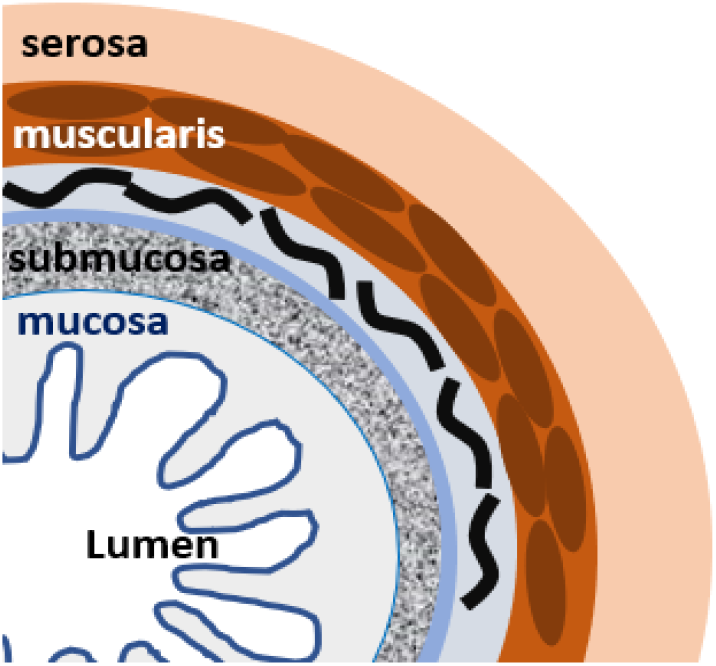
The cross section through the layers of the human colon.

The first *in-vivo* Raman measurements of human gastrointestinal tissue were published in 2000 by Shim et al. [8]. This study has shown that Raman spectroscopy can be successfully used for disease classification during *in-vivo* examination by using fiber probes coupled with Raman spectrometer. Raman studies of CRC have been published also by Andrade et al. [9]. Authors developed the diagnostic algorithm useful to establish the spectral differences of the complex colon tissues to find a characteristic Raman features of cancerogenesis. Popp group has performed the first CARS measurements of colon tissues [10]. The comparison with results obtained by using typical Stokes component of Raman spectra has shown many similarities, simultanously underlying the main advetage of CARS technqiue - the short acquisition time. Liu et al. used chemometrics methods: PCA and PLSDA to prove that Raman spectra can be effectivelly used to differentiate the normal and cancer human colon tisses [11]. In 2006, single living cells of the epithelium of CRC and control mucosa have been analyzed by Raman spectroscopy by Chen et al. [12]. PCA revealed a separation between epithelial cells of mucosa and cancerous tissue according to spectral signals assigned to nuclei and proteins with the sensitivity of 77.5% and the specificity of 81.3% [12].

In the last decade, several studies have shown that spectral histopathology (SHP) is capable of classifying different tissue types and especially diseased tissue such as cancer [13–17]. The advantage of spectroscopic techniques correlates with the fact that the measured vibrational spectra are integral signals of the proteome, genome, and metabolome. Other words, in one measurement is possible to obtain complex information about sample e.g. health or cancer status. Thus, when vibrational spectra are collected from distinct regions of tissue sections, variations in the spectral patterns can be detected and can be correlated with cancer areas [13–17].

Nowadays, early-stage cancer detection and edge detection of cancerous lesions is still challenging. The diagnostics is often based on invasive techniques whether a suspicious change - for example a colon polyp must be removed from the patient’s body. Generally, depending on the organ and the location of the suspicious tissue area, different biopsy methods can be used: punching out a cylinder of tissue (punch biopsy), aspiration of tissue or cells (fine-needle biopsy, fine-needle puncture), or sampling of tissue with a scalpel (excisional biopsy) or endoscopically with tiny forceps. Is necessary to highlight that these procedures are also time-consuming and expensive.

Also accurately detecting of cancer is a crucial and foremost step toward improving the survival rate of patients with colorectal cancer. Currently, colonoscopy and histopathology are standard screening and diagnostic techniques for colorectal tissues. Though colonoscopic screening has significantly increased the survival rate of patients with colorectal cancer, it remains a challenge to distinguish adenomas and early adenocarcinomas from benign hyperplasticpolyps using colonoscopy. Moreover, immunohistochemistry also has limitations due to the difficulty of analyzing large volumes of tissue sections by staining and the inability to detect multiple signals simultaneously. Also, the reproducibility and robustness of genomic data remain a concern due to the heterogeneity of tumors. It is, therefore, a real need to develop robust diagnostic and classification tools that have reproducibility and translational application with clinical samples. Raman microspectroscopy and imaging provide label-free identification and localization of cancer based on many fluorophores signals. Biomolecules such as proteins, lipids, or nucleic acids are Raman-active and thus provide molecular fingerprints that are highly sensitive and can reflect a specific tissue state or cellular phenotype. That’s why the development of new, spectroscopic cancer diagnostics methods in a form of SHP is so extremely valuable.

The other nowadays challenge is a cancer treatment. Some promising anti-cancer drugs are statins. Statins are understood to mean a group of organic, multifunctional chemical compounds. They occur both naturally and are made synthetically in laboratories. Depending on the structure of the statin compound, mevastatin, lovastatin, pravastatin, compactin (statins of natural origin), simvastatin (semi-synthetic statins), atorvastatin, rosuvastatin, pitavastatin, cerivastatin and fluvastatin (synthetic statins) can be distinguished. All these compounds have a pharmacophore group in their active form. Due to their pleiotropic effects, statins are also being used in oncology. In many experiments, both *in vitro* and *in vivo*, statins inhibited the proliferation of cancer cells, induced apoptosis, i.e. programmed cell death, and reduced the number of metastases or delayed their occurrence. They have also been observed to work synergistically with many drugs, not only with standard chemotherapeutics (cisplatin, doxorubicin), but also with preparations not used in the cancer treatment (bisphosphonates, saquinavir).

Generally, statins inhibit HMG-CoA reductase, the enzyme that converts HMG-CoA into mevalonic acid, a cholesterol precursor, but statins do more than just compete with the normal substrate in the enzymes active site. They alter the conformation of the enzyme when they bind to its active site. This prevents HMG-CoA reductase from attaining a functional structure. The change in conformation at the active site makes these drugs very effective and specific. Moreover, binding of statins to HMG-CoA reductase is reversible [18]. The inhibition of HMG-CoA reductase determines the reduction of intracellular cholesterol, inducing the activation of a protease which slices the sterol regulatory element binding proteins (SREBPs) from the endoplasmic reticulum. SREBPs are translocated at the level of the nucleus, where they increase the gene expression for LDL receptor. The reduction of cholesterol leads to the increase of LDL receptors, that determines the reduction of circulating LDL and of its precursors (intermediate density - IDL and very low density-VLDL lipoproteins) [19]. All statins reduce LDL cholesterol non-liniarly, dose-dependent, and after administration of a single daily dose. At least four mechanisms were proposed to explain statins antioxidant properties. (1) The hypocholesterolemic effect, resulting in reduced lipoprotein cholesterol, and thus, reduced level of oxidation substrate. (2) The decrease of cell oxygen production, by inhibiting the generation of superoxide by macrophages. Recently, it was demonstrated that statins can attenuate the formation of superoxide anion in endothelial cells, by preventing the prenylation of p21 Rac protein [20]. Statins can also prevent LDL oxidation by preserving the activity of the endogenous antioxidant system, like superoxide dismutase [21]. (3) The binding of statins to phospholipids on the surface of lipoproteins preventing the diffusion towards the lipoprotein core of free radicals generated during oxidative stress. (4) The potent antioxidative potential of the metabolites also results in lipoproteins protection from oxidation.

Because statins are structural analogs of 3-hydroxy-3-methyl-glutaryl-coenzyme A (HMG-CoA) their compete with it for the active site of HMGCoAR. As statins bind to the enzyme more strongly than its natural substrate, the reduction of HMG-CoA and the production of mevalonic acid (MVA) are inhibited [22,23]. Due to the fact that the cellular concentration of MVA depends on the activity of HMG-CoAR, and MVA is necessary for the subsequent reactions of the cholesterol synthesis pathway, this step is considered to be crucial for the whole process. For this reason, statins are used in the treatment of hypercholesterolaemia [22–27]. Moreover, statins increase the number of receptors for low-density lipoproteins on the surface of hepatocytes, which increases the absorption of cholesterol and additionally reduces its concentration in the blood [23,26,28,29]. Statins inhibit the progression of atherosclerosis and reduce the number of cardiovascular events in patients with ischemic heart disease (IHD) [30–32]. The beneficial effects of statin use in the treatment of IHD were also noted in patients with normal cholesterol levels, which suggests that statins also act in a mechanism independent of their cholesterol lowering effect [33]. Indeed, statins act on the cell and the body through several independent mechanisms. Due to their pleiotropic effect, the positive effects of their use are observed in the treatment of many diseases [34]. Statins have antiplatelet [35], antihypertensive [36,37] and anti-inflammatory properties [38,39]. Since the main indication for the use of statins is lipid disorders, which are a common disease, and this group of drugs is also used in other diseases, statins are among the most commonly prescribed drugs. Currently, there are reasons to use them also in the case of cancer.

The fact that mevalonate plays a key role in cell proliferation and that many malignant cells present an increased HMG-CoA reductase activity, suggests that a selective inhibition of this enzyme could lead also to a new chemotherapy for cancer disease. The obtained reduction of sterols synthesis by statins, suggests that inhibition of tumor cell growth can be related to the reduction of nonsteroidal isoprenoid compounds. The inhibitory effect on the synthesis of isoprenoid compounds formed in the side branches of the MVA pathway may play an important role in the anti-cancer properties of statins. These substances include dolichol, ubiquinone, isopentenes-loadenosine, geranylgeranyl pyrophosphate or farnesyl pyrophosphate [40]. Dolichol phosphate is also a carrier of extracellular sugar residues of proteoglycans, the effect of which can be associated with gene expression and with the change of antigenic properties of the cell membrane, intercellular interactions and the flow of information in signaling pathways [41]. The reduction in mevalonate synthesis also leads to a reduction in intracellular concentrations of farnesyl pyrophosphate (FPP) and geranylgeranyl (GGPP). These proteins are responsible for growth, differentiation, apoptosis, modulation of the actin cytoskeleton of cells, and thus for cell migration and adhesion. Mutant Ras or Rho proteins are typical for cancers [42].

In the presented studies the biochemical and nanomechanical characterization of human colon cells: normal CCD-18Co, cancer CaCo-2 and cancer CaCo-2 upon different statins supplementation will be analyzed by using Raman spectroscopy and imaging and AFM techniques. The influence of statins type will be also discussed.

## Materials and methods

### Cell lines and cell culture

CCD-18Co (ATCC^®^ CRL-1459™) and Caco-2 (ATCC^®^ HTB-37™) cell line was purchased from ATCC: The Global Bioresource Center. CCD-18Co cell line was cultured using ATCC-formulated Eagle’s Minimum Essential Medium with L-glutamine (catalog No. 30-2003). To make the complete growth medium, fetal bovine serum was added to a final concentration of 10%. Every 2-3 days, a new medium was used. The cells obtained from the patient are normal myofibroblasts in the colon. The biological safety of the CCD-18Co cell line has been classified by the American Biosafety Association (ABSA) as level 1 (BSL-1). The CaCo-2 cell line was also cultured according to the ATCC protocols. The CaCo-2 cell line was obtained from a patient - a 72-year-old Caucasian male diagnosed with colon adenocarcinoma. The biological safety of the obtained material was classified as level 1 (BSL1). To make the medium complete we based on Eagle’s Minimum Essential Medium with L-glutamine, with addition of a fetal bovine serum to a final concentration of 20%. The medium was renewed once or twice a week.

### Cultivation conditions

Cell lines (CCD-18Co, Caco-2) used in the experiments in this study were grown in flat-bottom culture flasks made of polystyrene with a cell growth surface of 75 cm^2^. Flasks containing cells were stored in an incubator providing environmental conditions at 37 °C, 5% CO_2_, 95% air.

### Raman Spectroscopy and Imaging

All maps and Raman spectra presented and discussed in this paper were recorded using the confocal microscope Alpha 300 RSA+ (WITec, Ulm, Germany) equipped with an Olympus microscope integrated with a fiber with 50 μm core diameter with a UHTS spectrometer (Ultra High Through Spectrometer) and a CCD Andor Newton DU970NUVB-353 camera operating in default mode at −60 °C in full vertical binning mode. 532 nm excitation laser line, which is the second harmonic of the Nd: YAG laser, was focused on the sample through a Nikon objective lens with magnification of 40x and a numerical aperture (NA = 1.0) intended for cell measurements performed by immersion in PBS. The average excitation power of the laser during the experiments was 10 mW, with an integration time of 0.5 s for Raman measurements for the high frequency region and 1.0 s for the low frequency region. An edge filter was used to filter out the Rayleigh scattered light. A piezoelectric table was applied to set the test sample in the right place by manipulating the XYZ positions and consequently record Raman images. Spectra were acquired with one acquisition per pixel and a diffraction grating of 1200 lines/mm. Cosmic rays were removed from each Raman spectrum (model: filter size: 2, dynamic factor: 10) and the Savitzky-Golay method was implemented for the smoothing procedure (order: 4, derivative: 0). All data was collected and processed using a special original software WITec Project Plus.

All imaging data were analyzed by Cluster Analysis (CA), which allows for grouping of a set of vibrational spectra that bear resemblance to each other. CA was executed using WITec Project Plus software with Centroid model and k-means algorithm, in which each cluster is represented by one vector of the mean. The normalization, model: divided by norm (divide the spectrum by the dataset norm) was performed by using Origin software according to the formula:

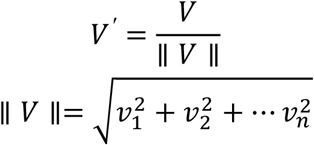

where: *v_n_* is the n^th^ V values.

The normalization was performed for all Raman spectra presented in the manuscript.

The Origin software was also used to perform ANOVA analysis necessary to indicate statistically significant results (means comparison: Tukey model, significance level: 0.05).

### AFM measurements

AFM measurements were performed by using PIK Instruments atomic force microscope with scanning range of 100 x 100 μm in the X and Y axes and 15 μm in the Z axis with a positioning resolution in the XY axis of 6 pm and in the Z axis of 0.9 pm, equipped with an inverted microscope, enabling measurements in air and liquid, in contact and tapping modes. Nanosurf C3000 software was used for AFM data collection. During measurements topography maps and nanomechanical properties of cells with and without supplementation of statins have been determined with the resolution 256×256 point per 60×60 μm. During measurements qp-Bio-AC-50 tips produced by Nanosensors with a spring constant 0.6 N/m were used. The analysis of AFM data was performed by using AtomicJ software [43] to obtain information about Young’s modulus of analysed biological samples.

For AFM measurements cells were cultured on Petri dishes filled with culture medium. Once the growing cells formed semi-confluent monolayer, the dish with cells was mounted on the AFM scanner, medium was replaced by PBS and the sample was measured within next 2–3 h (at room temperature and ambient conditions).

### Chemical compounds

Mevastatin (M2537-5MG), simvastatin (S6196-5MG), and lovastatin (PHR1285-1G), were purchased from Sigma-Aldrich, and used without additional purification. XTT proliferation Kit with catalogue Number 20-300-1000 was purchased from Biological Industries.

### XTT

In order to be able to perform clinical trials, a series of tests should be carried out to determine the activity of cells in terms of their metabolism and proliferation after exposure to specific substances. This is necessary because on this basis it is possible to determine whether a given chemical is producing a cytotoxic response. Initially, tests were developed to incorporate compounds such as 5-bromo-2-deoxyuridine (BrdU) or [H]-thymidine into the structure of DNA. Due to the inconvenience of this type of tests related to the need to use radioactive materials, expensive equipment or a time-consuming procedure, colorimetric methods have been developed. The basis of this method is the phenomenon observed for tetrazolium salts, which can be transformed by living cells as electron acceptors. As a result of this transformation, colored formazan compounds are formed. The first salt to be used in the colorimetric tests is 3-[4,5-dimethylthiazol-2-yl] −2,5-diphenyltetrazolium bromide known as the MTT salt. It is a positively charged compound, thanks to which it easily penetrates the cell, where it is reduced to a water-insoluble formazan compound. However, this method is also not perfect due to the need to dissolve the formazan compound crystals in an organic solvent. For this reason, a method was developed in which the MTT salt was replaced with the 2,3-bis [2-methoxy-4-nitro-5-sulfophenyl] −2H-tetrazoli-5-carboxanilide sodium salt, more widely known as the XTT salt. Unlike MTT, the XTT salt, when it enters the cell, is transformed into a product that can be dissolved in an aqueous medium. XTT, unlike the MTT salt, has a negative charge, so its permeability to the cell interior is low. This results in a reduction either at the cell surface or in the plasma membrane by the transmembrane electron transport chain.

One application of the XTT colorimetric assay is to test the viability of cells as a function of the compound that is active on them and the concentration of the compound. An example of this type of compound is statins. In the publication by Ludwig et al. the effect of three statins was investigated: atorvastatin, simvastatin and pravastatin [44]. For this purpose, a test was performed for each of the compounds for different concentrations of the tested substance. The statin compound was added after placing normal endothelial or cancer cells (CPAE) in a 96-well plate medium and after incubating the cells for 24 or 48h. In addition, a control was performed with only cells submerged in the pure culture medium. Then, after the addition of statins, the cells were incubated again for 4 h with formazan salts, after which it was possible to perform the measurement. Based on the obtained results, the survival curves of the studied cells were determined depending on the statin compound used.

### Determination of the appropriate statin concentration using the XTT test

For each cell type, XTT tests were performed 24 h and 48 h after the addition of mevastatin, lovastatin and simvastatin to the cells immersed in the culture medium. Preparation for the test included proper filling of the 96-well plate according to the procedure developed at the Institute of Applied Radiation Chemistry in Lodz. The wells were filled in such a way that each row contained a specific series of measurements. For example - in one row all plates were filled with medium, in another - control samples containing only cells immersed in the medium, and only in subsequent rows - cells in the medium with the addition of a specific concentration of selected statin. 6 different concentrations of each statin (mevastatin, lovastatin, simvastatin) were selected for the test: 1 μM, 5 μM, 10 μM, 25 μM, 50 μM, 100 μM. After completing each of the 96-well plates, the samples were incubated at 37 °C by 24 or 48 hours. After the time from the addition of statin, the XTT compound was added and the test was performed using the BioTek Synergy HT apparatus. The experiment was carried out after 3 hours from the addition of the reagent containing formazan salts. After the completion of the study, the obtained results had to be analyzed using a spreadsheet, resulting in a bar graph showing the effect of added statin concentration on the survival of the tested cell type, taking into account the time since the addition each of statin.

In our previous paper in which we investigated cancer human colon cells (CaCo-2), it was found that for cells the most appropriate concentration of mevastatin in the solution with medium would be 10 μM [45]. For each test, cell survivability at such a concentration was fluctuated in the range of 50-60%, which made it possible to conclude that at such a concentration, the effect of mevastatin on cells will be noticeable both in the study of nanomechanical and biochemical properties, and there will be enough living cells to allow will be conducting analyzes. Scheme 1 shows the results of XTT test obtained for Caco-2 human colon cancer cells supplemented with Lovastatin in various concentrations and in different time intervals.

Scheme 1 shows the results of XTT test obtained for Caco-2 human colon cancer cells supplemented with Lovastatin in various concentrations and in different time intervals.

**Scheme 1:**
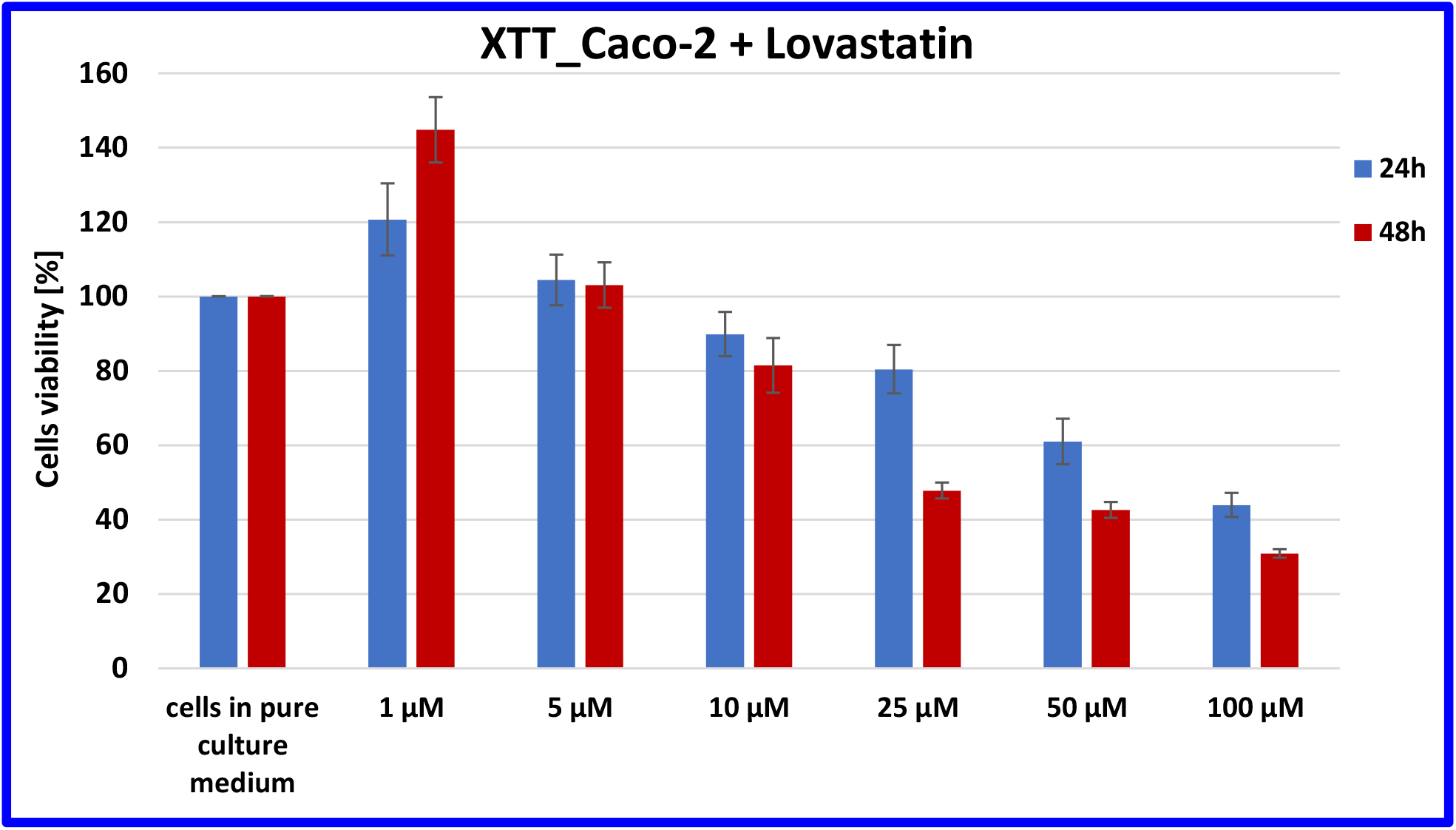
Results of XTT comparison of the percent viability for Caco-2 human colon cancer cells supplemented with different concentrations of Lovastatin in two different time intervals with the standard deviation ±SD.

As indicated by the obtained results, the most optimal concentration for the observation of the effect of Lovastatin supplementation is, similarly to the previous type of statin, 10 μM. Moreover, the concentration of the maintenance of a constant statin concentration of 10 μM enables reference to the comparison of the effect of other statins and to the previously published work describing the effect of Mevastatin, as we mentioned above [45].

Summarizing, taking into account the previously published results for Mevastatin, and presented data for Lovastatin we decided to use the same concentration for the other types of statins in order to compare their effect and compare the results to those obtained previously. The experimental idea applied in this way allows the most precise and unambiguous way to determine the effect of individual statins of the same concentration on the basic functions of the cell and to track possible biochemical changes.

In the presented studies, in the further part of the experiments (Raman spectroscopy and imaging and AFM), the effect of statins only on colon cancer cells (CaCo-2) after 24 and 48 hours was investigated because it is well known from the literature that statins are not destructive in normal cells, as is the case with most drugs used to treat cancer [46–51].

## Results and discussion

One of the main goals of our study was to determine the statistically significant differences between normal and cancer human colon cells including cancer cells supplemented by mevastatin, simvastatin and lovastatin based on vibrational features of them. Therefore to properly address these tasks we investigated systematically how the Raman imaging and Raman spectroscopy methods respond to *in vitro* normal and cancer human cells without and upon the supplementation by statins.

Herein, we present a valuable, fast and costless method for the cells structures visualization and the cells virtual staining, that adds the biochemical information given by the Raman intensity to the pseudo-colour images. These label-free images with high spatial resolution enable a direct analysis of all human colon cells substructures, which can help the tracking of biochemistry changes typical for cancerogenesis and can help the analysis of anti-cancer treatment.

Figure 1 shows the microscopy image, Raman image, Raman images of all cells substructures identify by using Cluster analysis algorithm, average Raman spectra typical for identified: lipid rich structures, mitochondria, nucleus, cytoplasm, cell membrane, and cell environment, and the average Raman spectra for the cell as a whole for human normal colon cells: CCD-18Co, human cancer colon cells CaCo-2, and human cancer colon cells CaCo-2 upon supplementation with simvastatin, lovastatin and mevastatin in 10 *μ*M concentration for 24h.

**Figure 1.**
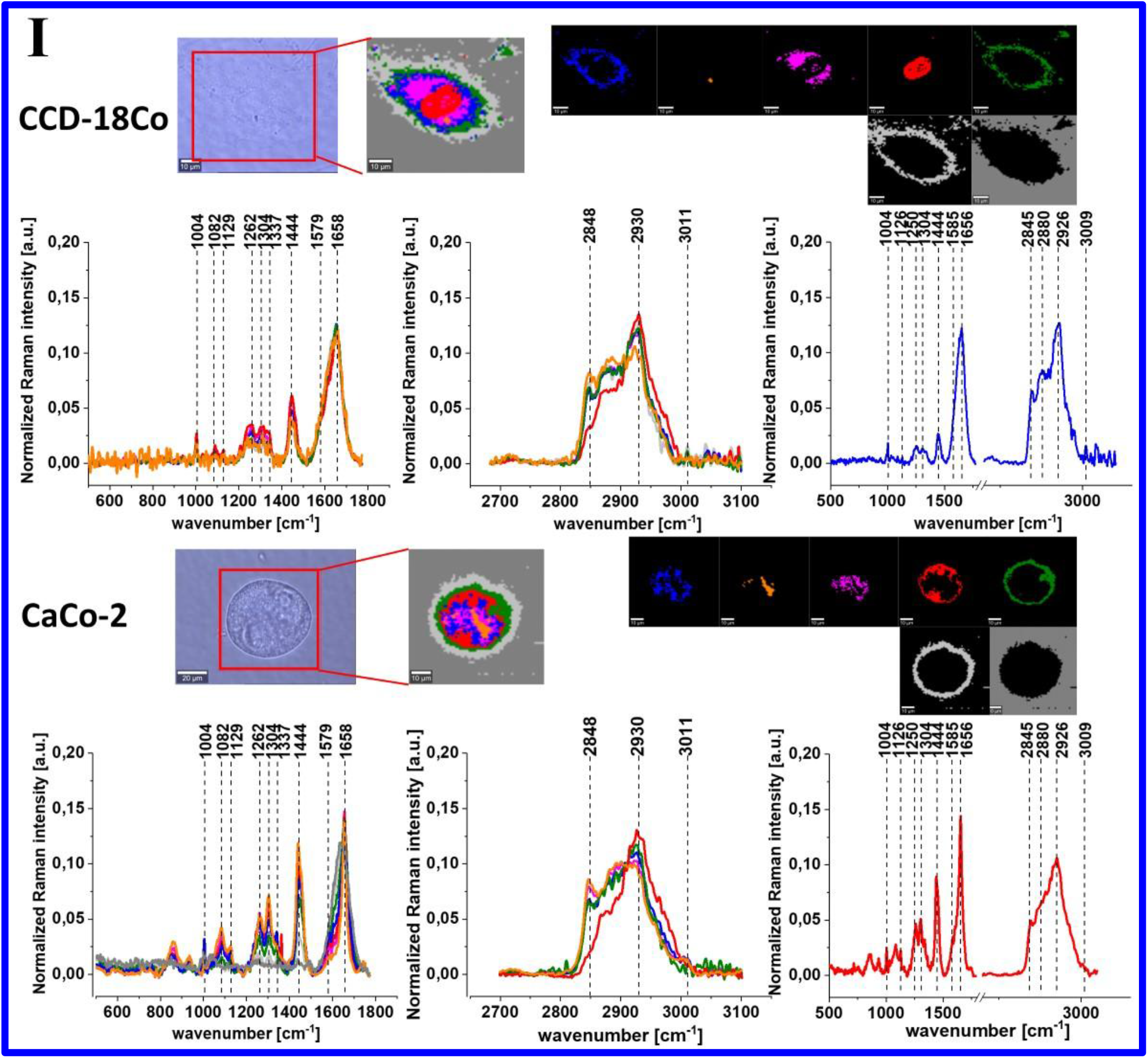
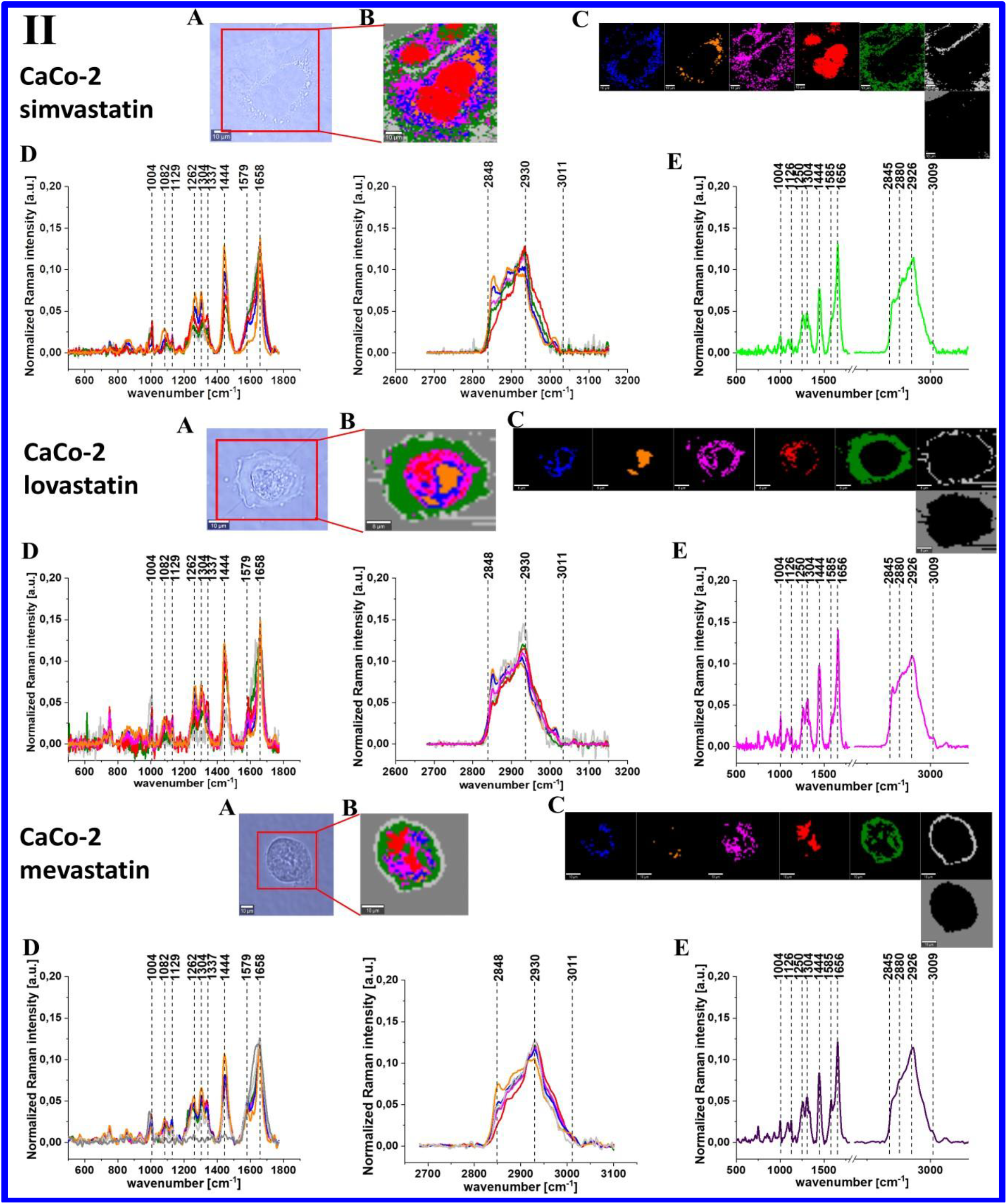
The microscopy image (A), Raman image (B) constructed based on Cluster Analysis (CA) method, Raman images of all clusters identified by CA assigned to: lipid-rich regions (blue and orange), mitochondria (magenta), nucleus (red), cytoplasm (green), cell membrane (light grey), and cell environment (dark grey) (C), the average Raman spectra typical for all identified clusters for low frequency and high frequency region (D), and the average Raman spectrum for cell as a whole (E) for human normal colon cells CCD-18Co(I), human cancer colon cells CaCo-2(I), and human cancer colon cells CaCo-2 upon supplementation with simvastatin, lovastatin and mevastatin in 10 μM concentration for 24h (II). All cells were measured in PBS, colors of the spectra correspond to the colors of clusters, excitation laser line 532 nm, the average Raman spectra calculated based on the data for 6 cells.

Generally, one can see from Figure 1 that based on the Raman spectra for each measurement the main biochemical components of single human colon cells can be identified. The fingerprint region of Raman spectra provides complex information on the biochemical composition of analyzed sample e.g. the peak at 755 cm^−1^ is associated with nucleic acids, DNA, tryptophan, nucleoproteins [52], the peaks c.a. 850 cm^−1^ can be assigned to tyrosine [40,52], a sharp peak at 1004 cm^−1^ corresponds to the aromatic amino acid phenylalanine [13–17,41], peak at 1126 cm^−1^ is typical for saturated fatty acids and cytochrome c, band at 1304 cm^−1^ corresponds to deformation vibration of lipids, adenine, cytosine [13–17,41], band at 1444/1452 cm^−1^ is typical for lipids and proteins, peak at 1585 cm^−1^ for CN_2_ scissoring and NH_2_ rock vibrations of mitochondria and phosphorylated proteins [13-17,41]. In the Raman spectra the peaks typical for proteins also can be observed in a form of the Amide I (C=O stretch) near 1656 cm^−1^, Amide II (N-H bend + C–N stretch) near 1557 cm^−1^, very weak and Amide III bands (C–N stretch + N–H bend) near 1260 cm^−1^ [13–17,41]. The high frequency peaks originates in the symmetric and antysymmetric stretching vibrations of C-H bonds found in lipids, glycogen, proteins, RNA, and DNA. Lipids and fatty acids including unsaturated fraction can be seen at 2845, 2880, 3009 cm^−1^. Proteins contribution, in the high frequency region, is observed at 2875, 2888, 2919 and 2926 cm^−1^.

Based on the Raman data obtained for normal, cancer and supplemented by statins cancer cells we can compare the vibrational futures of human colon cells using the average spectra calculated for cells as a whole and Raman band intensities ratios calculated for the main building blocks of biological samples: proteins, nucleic acids and lipids.

Figure 2 shows Raman band intensities ratios for selected Raman bands corresponding to nucleic acids 1004/1078, proteins 1004/1257, 1004/1658 and proteins and lipids 1004/1444 for four groups of human colon cells: normal human colon cells CCD18-Co: control group (labeled CCD-18Co, blue), cancer human colon cells CaCo-2 (labeled with CaCo-2, red), cancer human colon cells CaCo-2 incubated with statins in concentration of 10 *μ*M for 24 h (labeled with CaCo-2 simvastatin or lovastatin or mevastatin, 10 *μ*M, 24h, magenta), cancer human colon cells CaCo-2 incubated with statins in concentration of 10 *μ*M for 48 h (labeled with CaCo-2 simvastatin or lovastatin or mevastatin, 10 *μ*M, 48h, green) statistically significant data were marked with asterix (*).

**Figure 2.**
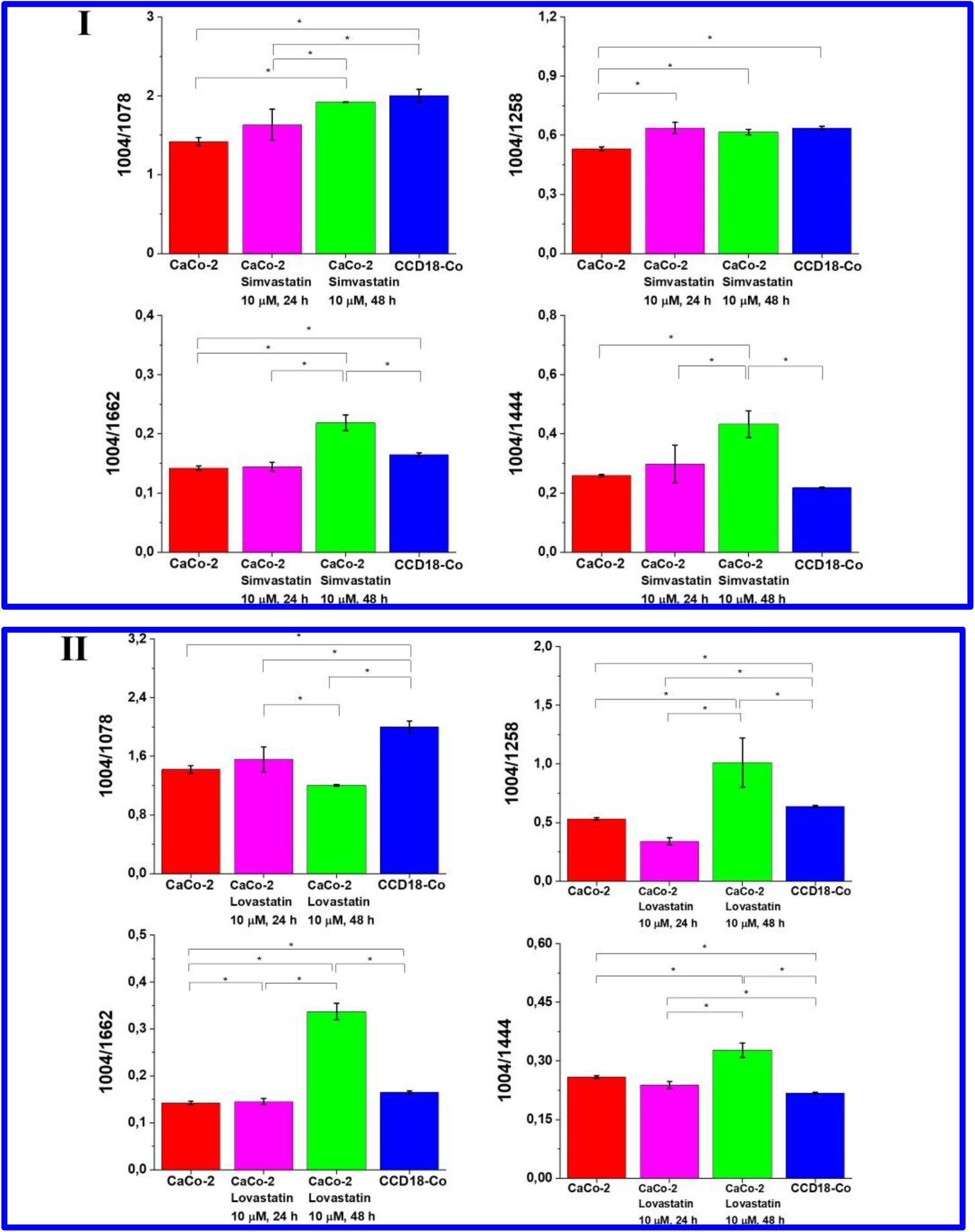
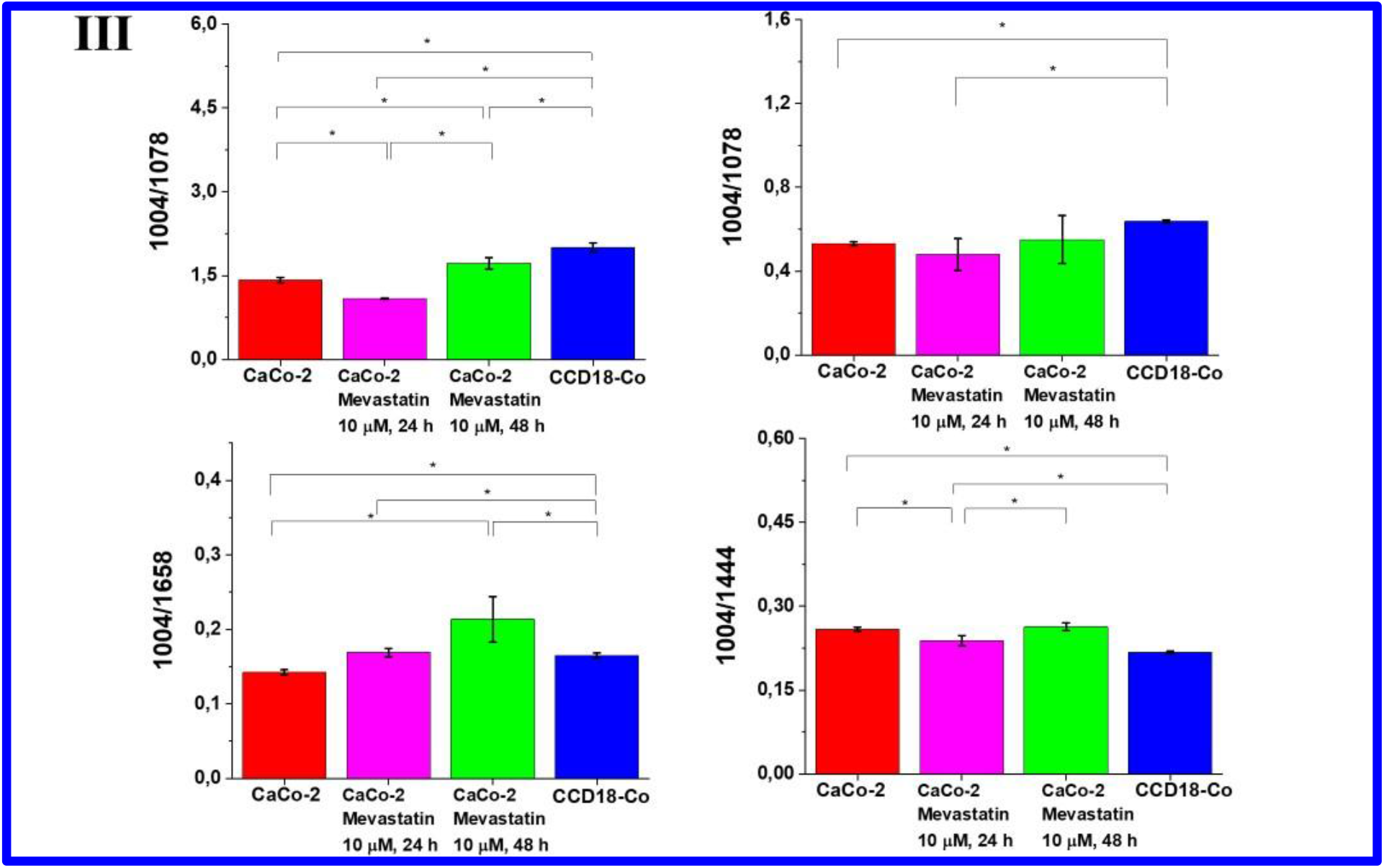
Raman band intensities ratios for selected Raman bands corresponding to nucleic acids 1004/1078, proteins 1004/1257, 1004/1658 and proteins and lipids 1004/1444 for four groups of human colon cells: normal human colon cells CCD18-Co: control group (labeled CCD-18Co, blue), cancer human colon cells CaCo-2 (labeled with CaCo-2, red), cancer human colon cells CaCo-2 incubated with statins in concentration of 10 μM for 24 h (labeled with CaCo-2 simvastatin (I) or lovastatin (II) or mevastatin (III), 10 μM, 24h, magenta), cancer human colon cells CaCo-2 incubated with statins in concentration of 10 μM for 48 h (labeled with CaCo-2 simvastatin (I) or lovastatin (II) or mevastatin (III), 10 μM, 48h, green) statistically significant data were marked with asterix (*).

One can see form Figure 2 that biochemical composition of normal, cancer and supplemented with statins cancer cells is different and the differences are observed for all the main chemical substituents: nucleic acids, proteins, lipids.

In the first step analyzing the differences between normal and cancer cells must be underlie that one of the factors responsible for the induction of cancer transformation are reactive oxygen species (ROS). The imbalance between the production of ROS and the efficiency of antioxidant systems leads to oxidative stress and, consequently, damage of DNA, proteins and lipids. Each day the content of the human colon can be described as a diverse mix of bile, mucus, gut microflora, fermentation products, unabsorbed food, and products of metabolism, including toxins, mutagens, and dissolved gases. In such an environment, the colon mucosa is constantly exposed to dietary oxidants and a variety of bacteria. Permanent exposure of the mucosa and the organism itself to unfavorable conditions may lead to uncontrolled oxidative stress and DNA damage, which may lead to the development of cancer disease.

Under the homeostasis conditions, ROS act as mediators and regulators of metabolism - they induce cell differentiation, activate many genes, including oncogenes, induce apoptosis, influencing the synthesis, release or inactivation of endothelial vasodilator factor (EDRF), have a dilating effect or contracting the wall of blood vessels, they increase the permeability of capillary walls, stimulate the transport of glucose to cells, serotonin to platelets. They influence the transmission of signals to cells and inside cells. They can become secondary transmitters both in the process of cell growth and death. It activates proteins that direct cell division (mitogenic activated protein). They take part in the body’s defense processes. Peroxides also regulate the synthesis of prostanoids.

Excessive production of ROS and depletion by the body’s antioxidant reserves is a phenomenon called “oxidative stress”. Oxidative stress leads to protein oxidation, which modifies their amount, structure and disrupts their function in the human body. Other main human body buildings components can also by damaged by ROS. The oxidation of lipids, damage to nucleic acids, depolymerization of hyaluronic acid and the accumulation of IgG can be observed. ROS also inactivate protease inhibitors, which increases the proteolytic effect of these tissue enzymes.

As was mentioned above the high concentrations of ROS trigger chain reactions, intensifying the processes of damaging biomolecules. Under ROS conditions e.g. the residues of polyunsaturated acids undergo oxidation and fatty acids, which are part of the phospholipids change cell membranes properties. Products of non-enzymatic peroxidation of lipids change the physical properties of cell membranes, which can lead to their damage.

Moreover, at the molecular level, ROS cause collagen degradation, disorders of the synthesis and inactivation of proteoglycans, enzyme inactivation, DNA strand breaks, the formation of guide mutations to cancer changes, inhibition of oxidative phosphorylation in mitochondria, structure disorders cytoskeleton (actin polymerization, disruption of microfiber laments), modification of antigenic properties cells and disturbance of intracellular calcium homeostasis.

Based on the data obtained by using Raman spectroscopy and imaging for proteins one can see in Figure 2 that amount of this class of compounds was different for normal and cancer human colon cells and was modulated by the adding of statins. In the presented analysis the intensity of the peak 1004 cm^−1^ was kept constant, which means that the decreasing of each ration correlates with the increasing of the amount of main building compounds of human colon cells: nucleic acids, proteins and lipids.

In Figure 2 for CaCo-2 cancer cells we observed the lower intensities for ratios 1004/1257 and 1004/1658 compared to CCD-18Co cells. Such a results were expected taking into account that the development of cancer is associated with the overexpression of proteins. However, for cancer cells incubated with statins in 10 μM concentration we noticed the statistically significant increase of analyzed ratios. This finding suggests that statins-induced inhibition of protein synthesis and the same protein dependent mechanism for cells death should be can underlined [53]. Protein synthesis is one of the most complicated biochemical processes undertaken by the cell, requiring approximately 150 different polypeptides and 70 different RNAs. In addition protein synthesis can be stopped when only a small fraction of the ribosome is inactivated by certain ribotoxins or when kinases associated with oxidative stress are activated [54]. The comparison between untreated human colon cells and cancer human colon cells upon statins supplementation shows that adding of statins effectively decreases the cell’s proteins level (the ratios 104/1258, 1004/1658 increase) especially for longer incubation time of 48h. One can see from Figure 2 that the strongest effect was observed for simvastatin.

Based on the data presented in Figure 2 for bands typical for nucleic acids one can notice that the intensity of the ratio typical for these compounds 1004/1078 decreases for CaCo-2 human cancer colon cells compared to control group – CCD-18Co corresponding to the normal human colon cells. This finding confirms that the synthesis of nucleic acids in cancer cells is enhanced, which is the expected result. Moreover, analyzing Figure 2 one can notice that the adding of statins modulates the amount of nucleic acids observed in CaCo-2 cancer cells. The ratio 1004/1078 increases confirming the reduction of DNA/RNA amount. Moreover, the concentration and incubation time dependence was observed. This finding is supported by scientific literature confirming, using traditional molecular biology methods, that decreased levels of DNA for cells interacting with statins is typical [55,56]. The strongest effect was observed for simvastatin.

The statistically significant differences between normal human colon cells, cancer human colon cells and cancer human colon cells upon statins supplementation have been found also for lipid components of analyzed samples. It is known that statins modulate the lipids composition of cells and tissue due to the influence on cholesterol level (in general, statins represent HMG-CoA reductase inhibitors, and are widely used for the treatment of hypercholesterolaemia) and the reduction of triglyceride concentrations. Results obtained based on intensity of Raman peaks related to lipids (peak at c.a. 1444 cm^−1^) confirmed that decreasing of intensity of peaks typical for lipids for cells treated by statins is observed and this effect is time and dose dependent [57]. The strongest effect was observed for simvastatin.

It is known from literature that lovastatin increases the concentration of cyclin inhibitors in the cell, arresting the G1 phase of the cancer cell cycle [58]. Due to the ability of statins to block proteasomal degradation of proteins [59,60], they show activity independent of the MVA pathway. The inhibitory effect of the proteasome, however, becomes apparent only at relatively high doses of statins. The activation of the peroxisome proliferator-activated receptor-g has been recently described as an additional anti-tumor mechanism of action for statins. It induces the production of tumor suppresor gene which is accompanied by a decrease in the phosphorylation of protein kinase B and mitogen activated protein kinases and the blockage of the cell cycle in the G1 phase [61].

A role in the cytostatic effect of statins can also be attributed to the induction of differentiation in cancer cells. In conclusion, the antiproliferative effect of statins has been confirmed in the treatment of gastric, pancreatic, breast, lung cancer, colon adenocarcinoma and acute myeloid leukemia [62–65]. Normal cells are also subject to this action. Statins inhibit the growth of normal endothelial cells, smooth muscles and fibroblasts [66,67]. However, the effect of statins on normal cells is much weaker, probably due to a lower proliferative potential and greater demand for its products in cancerous cells [40, 68–70]. The cytostatic effect of individual statins on different tumor cell lines is not identical. The effect of their use depends primarily on the dose and chemical properties, as well as the type of tumor.

Investigations regarding cells biochemistry were extended with analysis of nanomechanical properties of human colon cells: normal, cancer and cancer supplemented by statins. Figure 3 presents the data obtained during AFM measurements: topography maps, deflection maps, the topography maps in 3D visualization mode, and data for forward and backward traces measurements.

**Figure 3.**
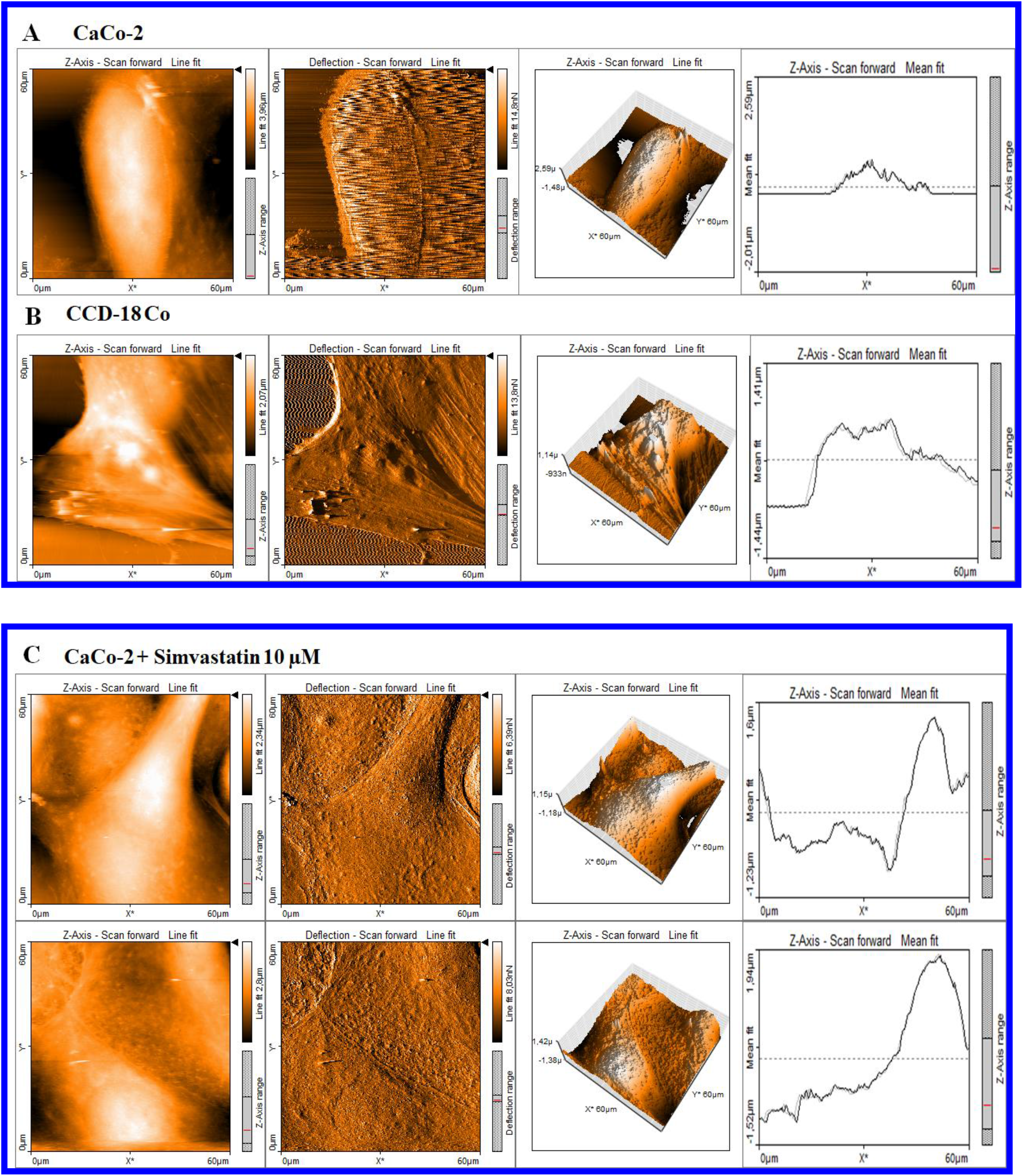
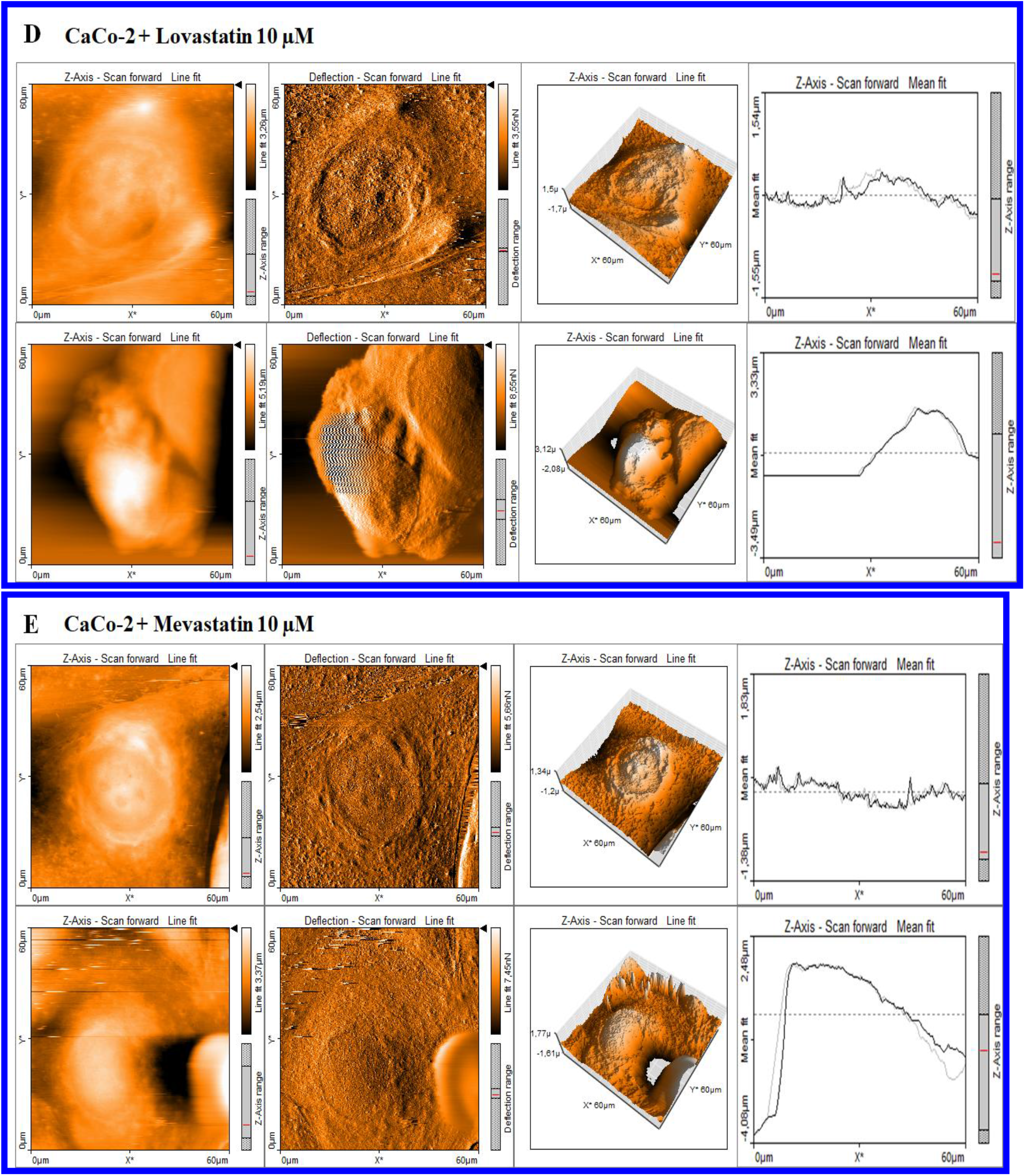
AFM topography maps of CaCo-2 (A), CCD-18Co (B), CaCo-2 supplemented by Semvistatin,10μM, 24h upper panel) and 48h (lower panel) (C), CaCo-2 supplemented by Lovastatin 10μM, 24h upper panel) and 48h (lower panel) (D), CaCo-2 supplemented by Mevastatin 10μM, 24h upper panel) and 48h (lower panel) (E) with deflection maps, 3D topography, and curves related to the topography measurements for forward and backward traces.

Figure 4 shows histograms related to the Young modulus calculated for each type of analyzed samples.

**Figure 4.**
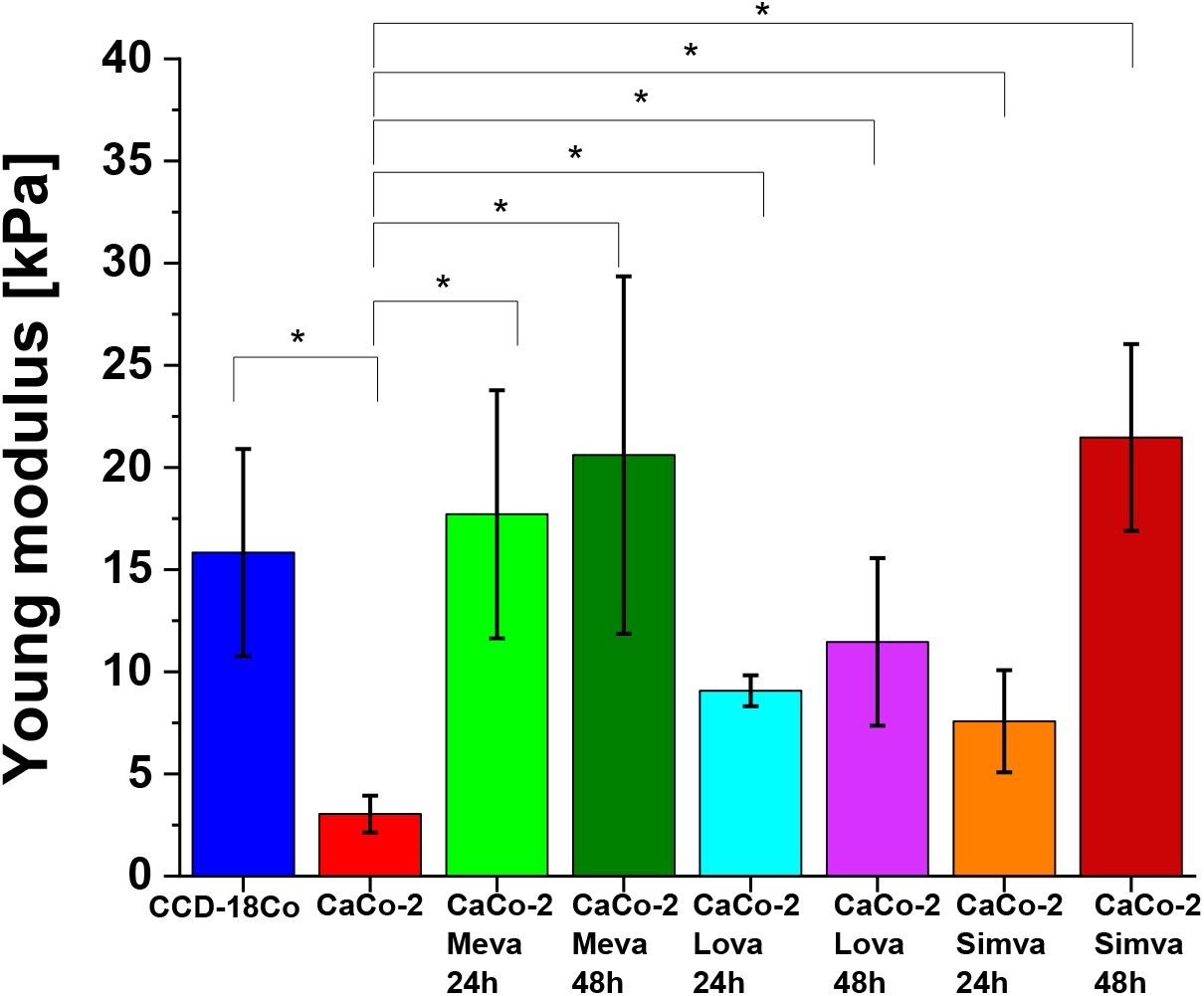
Young modulus values calculated for CCD-18Co (blue), CaCo-2 (red) CaCo-2 supplemented by: Mevastatin,10μM, 24h (light green), Mevastatin,10μM, 48h (dark green), Lovastatin,10μM, 24h (turquois), Lovastatin,10μM, 48h (violet), Simvastatin,10μM, 24h (orange), Simvastatin,10μM, 48h (brown).

One can see from Figure 4 that the cancer human colon cells CaCo-2 are more elastic compared to normal human colon cells CCD-18 Co and that the adding of statins in 10 μM concentration modulates the nanomechanical properties of cancers cells. The supplementation by using statins changed the elasticity of cancer cells and Young modulus values are more comparable to the elasticity of normal CCD-18 human colon cells. Based on ANOVA tests groups of CCD-18Co and CaCo-2 cells upon statins supplementation are not significantly different, while differences between CaCo-2 cells without and with statins supplementations are statistically significant. This finding confirms that the changes in skeleton organization of analyzed cells upon the statins supplementation occurred. The obtained result are consistent with literature data, which confirm higher flexibility of cancer cells compared to normal one [71].

Quantitatively results in Figure 4 proved that the value of Young’s Modulus for cancer cells is approximately 20% lower than for healthy cells. Supplementation with simvastatin causes a change in the value of the Young’s Modulus, and for 24 h supplementation it is a 2.5-fold increase in value, and for 48h supplementation - 7-fold in relation to cancer cells not subjected to supplementation. Supplementation with lovastatin also causes a change in the value of Young’s Modulus, and for 24 h supplementation it is a 3-fold increase in value, and for 48h supplementation 4-fold in relation to cancer cells not subjected to supplementation. Supplementation with mevastatin causes a change in the value of the Young’s Modulus, and for 24 h supplementation it is a 6-fold increase in value, and for 48h supplementation - 7-fold in relation to cancer cells not subjected to supplementation.

## Conclusions

The results proved that Raman imaging and spectroscopy are capable to differentiate human normal CCD-18Co and cancerous CaCo-2 colon cells and that vibrational spectra can be effectively used to efficient and accurate classification of single cells.

Based on the Raman spectra we visualized main substructures of single cells: nucleus, lipid structures, mitochondria, cytoplasm, cell membrane.

Atomic Force Microscopy allowed to characterize the nanomechanical properties of normal CCD-18Co and cancerous CaCo-2 human colon cells without and upon mevastatin supplementation.

The use of AFM to characterize elastic properties of normal and cancer cell lines justifies the idea of using nanomechanical parameters to track the changes typical for tumor development and antitumor treatment.

Accumulating evidence suggests that long-term use of lipophilic statins may also affect the overall incidence of cancer or the incidence of certain types of cancer. Moreover, statins may increase the sensitivity to chemotherapy and influence clinical outcomes in patients who have already been diagnosed with cancer.

*In vitro* studies have shown that statins inhibit tumor growth and induce apoptosis in colon cancer cell lines.

## Author Contributions

Conceptualization: BB-P; Funding acquisition: BB-P; Investigation: KB, BB-P; Methodology: BB-P, KB, Writing - original draft: KB, BB-P; Manuscript editing: KB, BB-P. All authors reviewed and provide feedback on the manuscripts. All authors have read and agreed to the published version of the manuscript.

## Acknowledgments

This article has been completed while the first author was the Doctoral Candidate in the Interdisciplinary Doctoral School at the Lodz University of Technology, Poland. Conflicts of Interest: The authors declare no competing interests. The funders had no role in the design of the study; in the collection, analyses, or interpretation of data; in the writing of the manuscript, or in the decision to publish the results.

